## Supplementary material for "InsectOR – webserver for sensitive identification of insect olfactory receptor genes from non-model genomes": S1-S4_Tables

### Application of InsectOR webserver on other insect species

**Table S1. InsectOR prediction of ORs in *Dufourea novaeangliae.*** (Known ORs: 112 [1])

| OR prediction method | InsectOR |
|---|---|
| No of predicted genes/gene-fragments | 107 |
| Proteins with 7tm_6 domain predictions | 95 |
| Complete hits with 7tm_6 | 80 |
| Partial hits with 7tm_6 | 15 |
| Normal hits with 7tm_6 | 86 |
| Pseudogene hits with 7tm_6 | 9 |

**Table S2. InsectOR prediction of ORs in *Apis florea.*** (Known ORs: 180 [2])

| OR prediction method | InsectOR |
|---|---|
| No of predicted genes/gene-fragments | 197 |
| Proteins with 7tm_6 domain predictions | 170 |
| Complete hits with 7tm_6 | 167 |
| Partial hits with 7tm_6 | 3 |
| Normal hits with 7tm_6 | 143 |
| Pseudogene hits with 7tm_6 | 27 |

**Table S3. InsectOR Prediction of ORs in *Anopheles gambiae.*** (Known ORs: 79 [3])

| OR prediction method | InsectOR |
|---|---|
| No of predicted genes/gene-fragments | 100 |
| Proteins with 7tm_6 domain predictions | 86 |

| | |
|---|---|
| Complete hits with 7tm_6 | 69 |
| Partial hits with 7tm_6 | 17 |
| Normal hits with 7tm_6 | 78 |
| Pseudogene hits with 7tm_6 | 8 |

**Table S4. InsectOR prediction of ORs in *Leptinotarsa decemlineata.*** (Known ORs: 37 [4])

| OR prediction method | InsectOR |
|---|---|
| No of predicted genes/gene-fragments | 84 |
| Proteins with 7tm_6 domain predictions | 54 |
| Complete hits with 7tm_6 | 12 |
| Partial hits with 7tm_6 | 42 |
| Normal hits with 7tm_6 | 48 |
| Pseudogene hits with 7tm_6 | 6 |
