## Supplementary material for "InsectOR – webserver for sensitive identification of insect olfactory receptor genes from non-model genomes": S1_File

```
# gffcompare v0.10.1 | Command line was:
#/home1/D/programs/gffcompare-0.10.1.Linux_x86_64/gffcompare -r
DmOrReference.gff -o DmOrperformanceComparison -p
DmOrconsensusNovelNotAllowed DmOrinsectOR.gff DmOrMaker.gff -Q -s
DmSeq.fasta -e 10
#

#= Summary for dataset: DmOrinsectOR.gff
#     Query mRNAs :      66 in      66 loci  (54 multi-exon transcripts)
#            (0 multi-transcript loci, ~1.0 transcripts per locus)
# Reference mRNAs :      66 in      61 loci  (65 multi-exon)
# Super-loci w/ reference transcripts:       58
#-----------------| Sensitivity | Precision  |
        Base level:    71.2     |    99.4    |
        Exon level:    33.6     |    43.9    |
      Intron level:    60.0     |    90.2    |
Intron chain level:    36.9     |    44.4    |
  Transcript level:    36.4     |    36.4    |
       Locus level:    39.3     |    36.4    |

     Matching intron chains:      24
       Matching transcripts:      24
              Matching loci:      24

          Missed exons:      58/247   ( 23.5%)
           Novel exons:       0/189   (  0.0%)
        Missed introns:      59/185   ( 31.9%)
         Novel introns:       2/123   (  1.6%)
           Missed loci:       1/61    (  1.6%)
            Novel loci:       0/66    (  0.0%)

#= Summary for dataset: DmOrMaker.gff
#     Query mRNAs :      25 in      25 loci  (24 multi-exon transcripts)
#            (0 multi-transcript loci, ~1.0 transcripts per locus)
# Reference mRNAs :      66 in      61 loci  (65 multi-exon)
# Super-loci w/ reference transcripts:       25
#-----------------| Sensitivity | Precision  |
        Base level:    38.4     |    83.8    |
        Exon level:    30.4     |    65.8    |
      Intron level:    38.4     |    79.8    |
Intron chain level:    23.1     |    62.5    |
  Transcript level:    22.7     |    60.0    |
       Locus level:    24.6     |    60.0    |

     Matching intron chains:      15
       Matching transcripts:      15
              Matching loci:      15

          Missed exons:     141/247   ( 57.1%)
```

```
           Novel exons:      11/114   (  9.6%)
        Missed introns:     102/185   ( 55.1%)
         Novel introns:      10/89    ( 11.2%)
           Missed loci:      34/61    ( 55.7%)
            Novel loci:       0/25    (  0.0%)

 Total union super-loci across all input datasets: 59
  (11 multi-transcript, ~1.3 transcripts per locus)
79 out of 79 consensus transcripts written in
DmOrperformanceComparison.combined.gtf (0 discarded as redundant)
```
