## Supplementary material for "InsectOR – webserver for sensitive identification of insect olfactory receptor genes from non-model genomes": S2_File

```
# gffcompare v0.10.1 | Command line was:
#/home1/D/programs/gffcompare-0.10.1.Linux_x86_64/gffcompare -r
referenceHlOr.gff -o HlOrperformanceComparison_NCBI7tm_6 -p
HlOrconsensusNovelNotAllowed insectORpredictedHlOr.gff
makerPredictedHlOr.gff ncbiGenes_7tm_6containing.gff -Q -s HlSeq.fasta -e
10
#

#= Summary for dataset: insectORpredictedHlOr.gff
#     Query mRNAs :     194 in     194 loci  (154 multi-exon transcripts)
#            (0 multi-transcript loci, ~1.0 transcripts per locus)
# Reference mRNAs :     151 in     150 loci  (140 multi-exon)
# Super-loci w/ reference transcripts:      147
#-----------------| Sensitivity | Precision  |
        Base level:    82.7     |    97.5    |
        Exon level:    53.3     |    67.8    |
      Intron level:    60.9     |    89.4    |
Intron chain level:    45.7     |    41.6    |
  Transcript level:    49.0     |    38.1    |
       Locus level:    49.3     |    38.1    |

     Matching intron chains:      64
       Matching transcripts:      74
              Matching loci:      74

          Missed exons:     182/868   ( 21.0%)
           Novel exons:       6/683   (  0.9%)
        Missed introns:     226/717   ( 31.5%)
         Novel introns:       4/489   (  0.8%)
           Missed loci:       2/150   (  1.3%)
            Novel loci:       0/194   (  0.0%)

#= Summary for dataset: makerPredictedHlOr.gff
#     Query mRNAs :     114 in     114 loci  (106 multi-exon transcripts)
#            (0 multi-transcript loci, ~1.0 transcripts per locus)
# Reference mRNAs :     151 in     150 loci  (140 multi-exon)
# Super-loci w/ reference transcripts:       89
#-----------------| Sensitivity | Precision  |
        Base level:    73.3     |    84.9    |
        Exon level:    31.0     |    39.6    |
      Intron level:    37.5     |    47.6    |
Intron chain level:     3.6     |     4.7    |
  Transcript level:     4.0     |     5.3    |
       Locus level:     4.0     |     5.3    |

     Matching intron chains:       5
       Matching transcripts:       6
              Matching loci:       6
```

```
          Missed exons:     280/868   ( 32.3%)
           Novel exons:     109/679   ( 16.1%)
        Missed introns:     207/717   ( 28.9%)
         Novel introns:      68/565   ( 12.0%)
           Missed loci:      23/150   ( 15.3%)
            Novel loci:       0/114   (  0.0%)

#= Summary for dataset: ncbiGenes_7tm_6containing.gff
#     Query mRNAs :      60 in      60 loci  (55 multi-exon transcripts)
#            (0 multi-transcript loci, ~1.0 transcripts per locus)
# Reference mRNAs :     151 in     150 loci  (140 multi-exon)
# Super-loci w/ reference transcripts:       57
#-----------------| Sensitivity | Precision  |
        Base level:    54.3     |    80.1    |
        Exon level:    32.7     |    54.7    |
      Intron level:    38.2     |    59.7    |
Intron chain level:     8.6     |    21.8    |
  Transcript level:     9.3     |    23.3    |
       Locus level:     9.3     |    23.3    |

     Matching intron chains:      12
       Matching transcripts:      14
              Matching loci:      14

          Missed exons:     428/868   ( 49.3%)
           Novel exons:      80/519   ( 15.4%)
        Missed introns:     145/717   ( 20.2%)
         Novel introns:      64/459   ( 13.9%)
           Missed loci:      21/150   ( 14.0%)
            Novel loci:       0/60    (  0.0%)

 Total union super-loci across all input datasets: 108
  (57 multi-transcript, ~3.5 transcripts per locus)
378 out of 378 consensus transcripts written in
HlOrperformanceComparison_NCBI7tm_6.combined.gtf (0 discarded as
redundant)
```
